# Mouse Genomic Associations with *Ex Vivo* Sensitivity to Simulated Space Radiation

**DOI:** 10.1101/2022.03.03.482929

**Authors:** Egle Cekanaviciute, Duc Tran, Hung Nguyen, Alejandra Lopez Macha, Eloise Pariset, Sasha Langley, Giulia Babbi, Sherina Malkani, Sébastien Penninckx, Jonathan C. Schisler, Tin Nguyen, Gary H. Karpen, Sylvain. V. Costes

## Abstract

Exposure to ionizing radiation is considered by NASA to be a major health hazard for deep space exploration missions. Ionizing radiation sensitivity is modulated by both genomic and environmental factors. Understanding their contributions is crucial for designing experiments in model organisms, evaluating the risk of deep space (i.e. high-linear energy transfer, or LET, particle) radiation exposure in astronauts, and also selecting therapeutic irradiation regimes for cancer patients. We identified single nucleotide polymorphisms in 15 strains of mice, including 10 collaborative cross model strains and 5 founder strains, associated with spontaneous and ionizing radiation-induced *ex vivo* DNA damage quantified based on immunofluorescent 53BP1^+^ nuclear foci. Statistical analysis suggested an association with pathways primarily related to cellular signaling, metabolism, tumorigenesis and nervous system damage. We observed different genomic associations in early (4 and 8 hour) responses to different LET radiation, while later (24 hour) DNA damage responses showed a stronger overlap across all LETs. Furthermore, a subset of pathways was associated with spontaneous DNA damage, suggesting 53BP1^+^ foci as a potential biomarker for DNA integrity in mouse models. Based on our results, we suggest several mouse strains as new models to further study the impact of ionizing radiation and validate the identified genetic loci. We also highlight the importance of future human *ex vivo* studies to refine the association of genes and pathways with the DNA damage response to ionizing radiation and identify targets for space travel countermeasures.

## Introduction

Interest in human deep space exploration has increased significantly over the last decade, especially focusing on detrimental health effects caused by ionizing radiation beyond the protective magnetic field of the Earth (Durante and Cucinotta 2008). For example, a Mars mission estimated to last nearly three years will expose astronauts to radiation doses of up to 1 Sv, which is nearly 300 times higher than a computer tomography scan (Cucinotta and Durante 2006). The most biologically concerning source of ionizing radiation beyond the Earth’s magnetosphere is galactic cosmic rays (GCRs), composed of 90% energetic protons, 9%^4^He nuclei and 1% high mass-high charge (HZE) particles. HZE particles are ions between ^4^He and ^56^Fe that induce significant biological damage due to their high linear energy transfer (LET), and are only partially mitigated by protective shielding (Cucinotta and Durante 2006; Durante and Cucinotta 2008). Therefore, understanding the biological effects of HZE particles is essential for identifying biomarkers for individual sensitivity and resistance and developing countermeasures to reduce astronaut health risks.

One of the main adverse effects of exposure to ionizing radiation, especially HZE particles, is DNA damage (Cardis *et al*. 1995; National Research 2006; Cardis *et al*. 2007), which leads to carcinogenesis, central nervous system impairments and immune dysregulation (Gasser *et al*. 2005; Braganza *et al*. 2012; Wang *et al*. 2016). An early mouse experiment has shown that HZE particles have a relative biological effectiveness (RBE) as high as 30 for causing Harderian gland cancer (Alpen *et al*. 1994), however, the mouse strain is not reported, making it unable to infer whether the effect was associated with strain-dependent radiosentivity.

Although the carcinogenic and DNA damage-related effects of HZE particles in animal models and human cells are the focus of numerous studies, the genomic and environmental factors that contribute to *individual* sensitivity among subjects remain elusive (Barcellos-Hoff *et al*. 2015; Cucinotta *et al*. 2017). Meanwhile, the known genetic predispositions to certain cancers (Wicking *et al*. 1997; Ariffin *et al*. 2014), demonstrate the importance of incorporating personalized genetic backgrounds when assessing carcinogenic and other risks following exposure to ionizing radiation. Thus, understanding the genes and pathways linked with HZE particle sensitivity may suggest mechanisms to be targeted for developing countermeasures to avoid the radiation hazards during prolonged space travel.

Currently the health risk from ionizing radiation exposure is typically estimated using health records of survivors of the atomic bombs in Japan and nuclear reactor workers (Cardis *et al*. 1995; Preston *et al*. 2003a; Preston *et al*. 2003b; Preston *et al*. 2004; Cardis *et al*. 2007; Preston *et al*. 2007), who have almost exclusively been exposed to low-LET gamma radiation. However, the translatability of health effects induced by low-LET radiation to predictions of high-LET radiation-induced risks is limited (Cucinotta 2015), because some forms of physiological damage, for example, impairments in angiogenesis and immune responses, are unique to high-LET radiation (Paul *et al*. 2020; Wuu *et al*. 2020). As a result, model organisms are essential tools to understand space-relevant radiation responses *in vivo* (Barcellos-Hoff *et al*. 2015).

In particular, mice are commonly used to model genotype-phenotype associations in human diseases due to high similarities between genomes (Mouse Genome Sequencing *et al*. 2002). A recent study used genetically diverse HS/Npt stock mice to characterize high-LET particle-induced tumor formation *in vivo* (Edmondson *et al*. 2020) and compared them to spontaneous and gamma-ray-induced tumors. The tumor histotypes developed by the mice were strongly heritable, however, neither high-LET particles nor low-LET gamma radiation were shown to have a specific histotype signature. Thus, we may conclude that there is low correlation between low and high LET responses across the population, but high correlation between the responses of an individual with the same genetic background to both low and high LET radiation. Therefore, it is important to account for genetic background when attempting to predict the high LET responses from low LET responses.

In order to identify mouse genes associated with space radiation risks, we used the *collaborative cross* (CC) mice (Churchill *et al*. 2004), a panel of recombinant inbred strains that captures 90% of the known variation among laboratory mice. This system allows high-resolution mapping of genomic associations with different phenotypes, including diseases and reactions to various environmental stressors, due to their genomic diversity and long-term genetic stability. The U.S. Department of Energy Low Dose Scientific Focus Area utilizes CC mouse model due to the high levels of genetic variation distributed randomly across the strain genomes (Snijders *et al*. 2016). This model enhances the probability of genomic associations to phenotypes using a relatively small number of strains and permits other investigators to study the phenotypes of interest further by replicating the same specific, individual genotypes of mouse strains.

In this work, we utilized the CC model to identify genetic associations with *ex vivo* DNA damage responses, We and others have previously reported that biomarkers labeling DNA double strand breaks are suitable surrogate markers of sensitivity to ionizing radiation (Rube *et al*. 2008; Ochola *et al*. 2019; Penninckx *et al*. 2019; Pariset *et al*. 2020b; Penninckx *et al*. 2020). More recently, our group used *ex vivo* samples from a combination of 15 mouse strains (5 inbred and 10 CC) to demonstrate that both their baseline levels of spontaneous DNA damage and their dose responses to ionizing radiation were highly variable among different strains based on quantification of immunofluorescent foci of tumor suppressor p53-binding protein 1 (53BP1) (Penninckx *et al*. 2019; Pariset *et al*. 2020b), which is a key component of DNA double-stranded break repair. Here we followed up on this result by conducting an exploratory genome-wide association study using the same 15 mouse strains to identify SNPs and pathways associated with spontaneous DNA damage, as well as DNA damage responses to different types of ionizing radiation. Specifically, we compared the responses to low-LET X-rays and two types of HZE particle irradiation (350 MeV/n ^40^Ar at 104 keV/μm LET; and 600 MeV/n ^56^Fe at 170 keV/μm LET) at different doses and post-irradiation time points. We further evaluated the pathways associated with responses to HZE particles due to their relevance to health risks during deep space exploration.

By identifying candidate genes and pathways related to space health risks, we provide new directions for selecting the appropriate mouse models for space biology and space radiation studies: different mouse strains might be more suitable for analyzing high-vs. low-LET responses and different aspects of DNA damage and repair kinetics. Our results also suggest targets for evaluating and mitigating DNA damage caused by deep space radiation at the level of an individual subject, opening the door for precision space medicine. All raw data and associated genotypes for each animal are available in the Open Science NASA platform GeneLab, a life science database focused on spaceflight-relevant experiments with the highest standards for rich metadata (Berrios *et al*. 2021) and curated radiation dosimetry information (Beheshti *et al*. 2018), which are essential features to enable further mining of these data by the scientific community.

## Materials and Methods

### Mouse strains

Collaborative Cross (CC) mice are inbred mice designed by the Jackson Laboratory (Bar Harbor, ME) and developed at University of North Carolina, Chapel Hill (Srivastava *et al*. 2017) that are derived from five classic inbred strains (A/J, C57BL/6, 129S1/SvImJ, NOD/ShiltJ and NZO/H1LtJ) and three wild-derived sub-strains (CAST/EiJ, PWK/PhJ and WSB/EiJ). 10 mouse strains from the CC panel - CC002, CC011, CC013, CC019, CC032, CC037, CC040, CC042, CC051 and CC061 - were chosen for our experiment. In addition to CC mice strains, five inbred reference strains: C57BL/6J, BALB/cByJ, B6C3F, C3H/HeMsNsrf and CBA/CaJ were also included in this study. In total, 72 animals were used, with 37 females and 35 males.

### Irradiation

Detailed experimental methods for cell extraction and irradiation are described in our previous work (Penninckx *et al*. 2019). All experiments were performed following IACUC protocol no. 306002; Lawrence Berkeley National Laboratory (LBNL), Berkeley, CA. X-ray experiments were conducted at LBNL using a 160-kVp Faxitron X-ray machine (Lincolnshire, IL), while particle irradiation experiments were conducted at NASA Space Radiation Laboratory (NSRL) beam line at Brookhaven National Laboratory (Upton, NY).

Briefly, skin fibroblasts from ear punches were collected from mice at 10-12 weeks of age, grown as primary cell cultures and frozen at different passages. Cells were thawed and grown for 24 – 48 hours before being irradiated with X-rays or two different HZE particles: 350 MeV/n ^40^Ar and 600 MeV/n ^56^Fe, which have 104 and 170 keV/μm LET, respectively. At the time of irradiation, fibroblasts were 80% confluent for ^40^Ar and 90% confluent for ^56^Fe. For particle irradiations, two fluences (1.1 and 3 particles/100 μm^2^) were used, corresponding to four distinct doses depending on their respective LET (0.30 and 0.82 Gy for 600 MeV/n ^56^Fe; 0.18 and 0.50 Gy for 350 MeV/n ^40^Ar). For X-ray irradiation, 0.1 and 1 Gy doses were used instead. The dose rate for both X-ray and particle irradiations was 1 Gy/min.

### Immunocytochemistry

At 4, 24 and 48 hours post-irradiation, with an additional time point at 8 hours for 600 MeV/n ^56^Fe particle irradiations, cells were fixed with 4% paraformaldehyde (Electron Microscopy Sciences, Hatfield, PA) in phosphate buffered saline (PBS) for 15 minutes at room temperature. After the incubation period, cells were washed with PBS 3 times for a total of 5 minutes each. Plates were then filled with PBS (300 μl/well), sealed and kept at 4°C until immunostaining. All experiments were performed in duplicate.

For immunostaining 53BP1 to quantify DNA double strand breaks, cells were washed with PBS, permeabilized with 0.1% Triton-X for 20 minutes and blocked with 3% BSA in PBS for 1 hour. Cells were then incubated in a rabbit polyclonal anti-53BP1 primary antibody (Bethyl Labs #IHC-00001) at 1:400 in blocking buffer (3% BSA in PBS) for 1 hour, followed by 2 washes in 0.1% Tween-20 in PBS and a subsequent incubation with Alexa Fluor 488 goat anti-rabbit secondary antibody (ThermoFisher Scientific #A11034) at 1:400 in blocking buffer for 1 hour, followed by 2 washes in 0.1% Tween-20 in PBS. Nuclear staining was performed with DAPI at 1:1000 in PBS for 5 minutes (ThermoFisher Scientific, #D1306), followed by 2 final washes in 0.1% Tween-20 in PBS and resuspension in PBS.

### Imaging

After staining, cells were imaged and quantified using a high-throughput semi-automated microscope developed in-house by our group (40X 0.95 NA dry lens, Nikon), and images were analyzed using an automated nuclear foci quantification MATLAB®algorithm previously published by our group (Penninckx *et al*. 2019; Pariset *et al*. 2020a). The foci quantification was set at a target of at least 800 cells per sample based on previous publications (Costes *et al*. 2007). All raw data are available online (see Data Availability), while images and quantification algorithm are available upon request.

### Genotyping

Genomic DNA was extracted from fibroblasts obtained from each individual mouse from each of the five non-CC inbred strains using AllPrep DNA/RNA Mini kit (Qiagen, #80204). The concentration and purity of extracted DNA was determined using a NanoDrop 2000 UV-vis spectrophotometer (ThermoFisher Scientific) based on 260 nm/280 nm and 260 nm/230 nm ratios. All DNA samples contained a minimal concentration of 20 ng/μL and were shipped to GeneSeek (Neogen Genomics, Lincoln, NE, USA). Single nucleotide polymorphism (SNP) analysis was conducted using MegaMouse Universal Genotyping Array (MegaMUGA platform). The original MUGA array was developed on Illumina Infinium platform in cooperation with Neogen Inc and contained 7851 SNP markers all spaced uniformly approximately ~325 kb across the mouse reference genome. The MegaMUGA SNP array that was used provided a 10-fold higher marker density than MUGA (77,808 markers) and additional 14,000 probes detecting variants segregating in wild-derived strains. Collaborative cross genotypes were obtained from the GigaMUGA database (Morgan *et al*. 2015). Only the overlapping SNPs that were found in both platforms were used for all analysis.

## Statistical Analysis

### Radiosensitivity phenotypes

As published previously (Penninckx *et al*. 2019; Pariset *et al*. 2020b), pontaneous DNA damage (Background; BGD) phenotype for each mouse strain was defined as the average number of 53BP1^+^ foci per nucleus in the control group (0 Gy sham irradiation), assessed at 4 different time points: 4, 8, 24 and 48 h for all three radiation qualities, averaged across all analyses after removing statistical outlier values (2 standard deviations above or below the mean for each strain).

Radiation-induced DNA damage (Foci per Gray; FPG) phenotype was defined as the average increase in 53BP1^+^ foci per nucleus, per 1 Gy of irradiation. The FPG phenotype was quantified in response to irradiation by X-rays and HZE particles (350 MeV/n ^40^Ar, 600 MeV/n ^56^Fe), averaged per mouse strain, separately for each radiation quality and time point.

### Genotype-phenotype associations

The significance of SNP-phenotype associations was determined using the R *qtl2* package (Broman *et al*. 2019) that includes kinship information between samples. For FPG calculation in response to each radiation quality, time points were used as covariates. The following linear model was used to infer the relationship between phenotype and genotype:

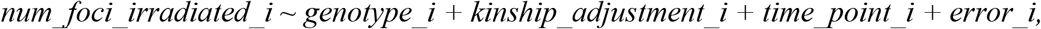

where *i* indexes the mouse.

The output of this calculation is the LOD (limit of detection) score for each SNP. LOD scores can be converted to p-value using:

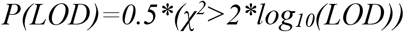

where 2*log_10_(LOD) follows the chi-square distribution with degree of freedom = 1 (Nyholt 2000).

### SNP annotation and pathway analysis

**Supplementary Figure 1** shows the overall workflow of pathway analysis. The *p* values for each SNP were calculated using *qtl2* R package and serve as an input of the workflow. Bioconductor package *BiomaRt v.105* (Durinck et al. 2005; Durinck et al. 2009) was used to get gene annotations and positions, mapping SNPs within 25,000 bp of a gene. *Reactome v.78* (Jassal et al. 2020) *v*. was used to get pathway information.

For each gene, the p-values of the SNPs were merged using two different methods: Fisher and the additive method addCLT (Nguyen *et al*. 2016). The Fisher method produces a list of *p* values, one value per gene. These *p* values are subsequently used either to create a pre-ranked gene list using FGSEA (Korotkevich *et al*. 2021) or correct them using False Discovery Rate (FDR) with a threshold of 0.05 to select differentially expressed genes for hypergeometric test (ORA), and both FGSEA and ORA are used for pathway analysis. Thus, our analysis results in 4 lists of pathway *p* values, generated by the following combinations of methods: Fisher/FGSEA, Fisher/ORA, addCLT/FGSEA, addCLT/ORA.

For each pathway, these 4 *p* values are combined using addCLT to produce a single additive *p* value per pathway. The additive p-values of all pathways are then corrected using FDR. The final pathways with FDR-corrected p-values <0.05 are considered significantly associated with the phenotype (either BGD, or FPG for each radiation type and time point). Our main motivations behind combining *p* values at the end of the pipeline are a) accounting for multiple sources of weak but consistent evidence and b) combining the evidence generated by different methods to prevent the flaws of any individual method from compromising the results. All pathway graphs were created using R packages *ggplot2*, *ggraph*, *igraph* and *UpSetR*.

All code used for the analysis is available in **Supplementary Materials** (see **Data and Code Availability** section).

## Results and Discussion

### Experimental approach

Our study aimed to identify the genetic differences and biological mechanisms responsible for the variability of DNA damage responses to ionizing radiation established in previous investigations of 15 mouse strains (5 inbred, 10 CC) that had been established in previous investigations (Penninckx *et al*. 2019). DNA damage response patterns were defined by quantifying 53BP1^+^ radiation-induced foci between 4 h and 48 h post-irradiation in mouse skin fibroblasts irradiated *ex vivo*. We used two doses of low-LET X-rays (0.1 and 1 Gy), and two fluences of high-LET 350 MeV/n ^40^Ar and 600 MeV/n ^56^Fe ions (1.1 and 3 particles/100μm^2^, which respectively correspond to 0.18 and 0.5 Gy for 350 MeV/n ^40^Ar, and 0.3 and 0.82 Gy for 600 MeV/n ^56^Fe). The experimental design is depicted in **Fig. 1A**.

**Figure 1.**
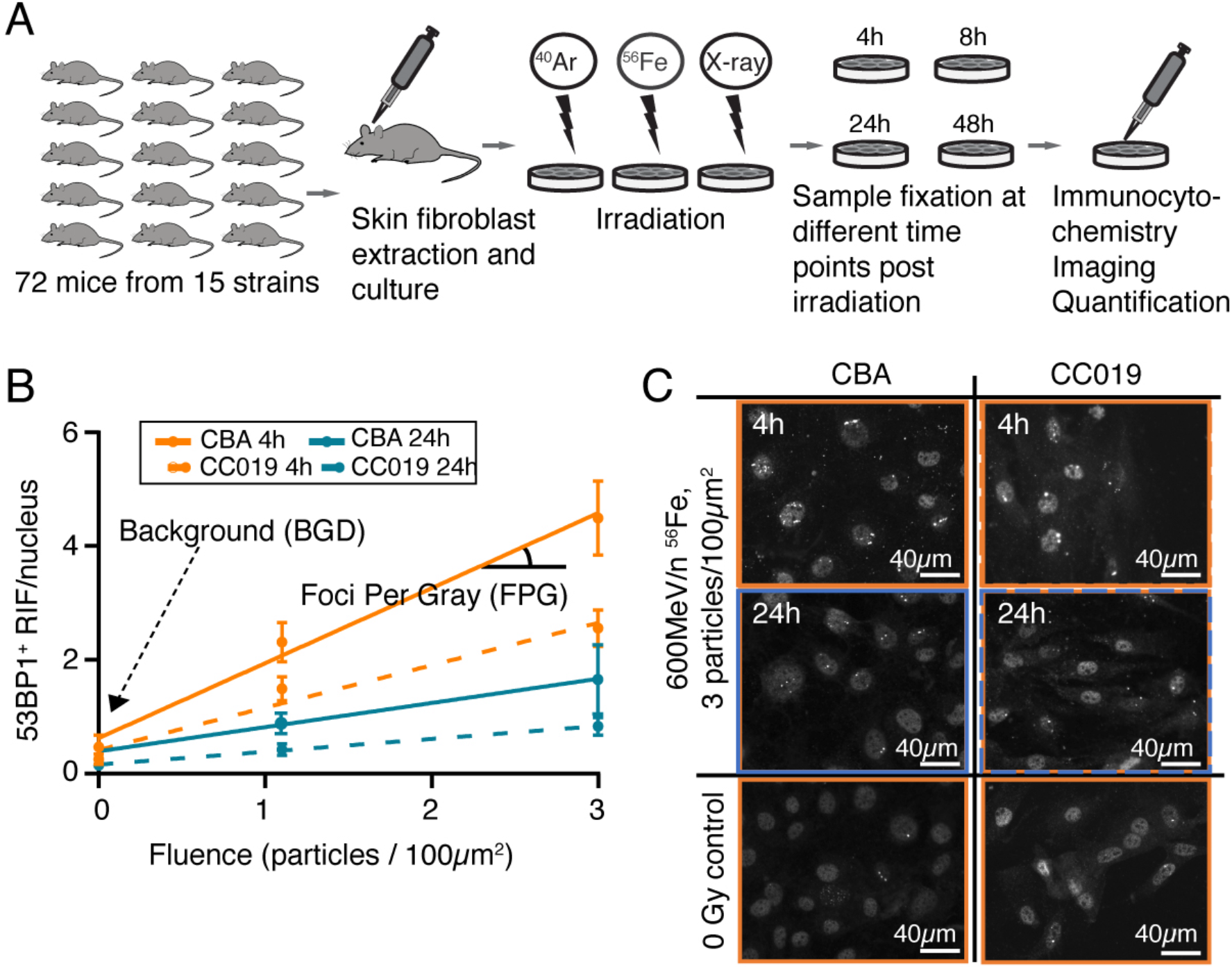
Quantification of radiation-induced DNA damage in 15 mouse strains. **A**. Experimental design. **B**. Representative graph showing Foci Per Gray (FPG) and Background (BGD) phenotypes in two mouse strains (CBA and CC019) at two time points (4 h and 24 h) post-irradiation as a function of 600 MeV/n ^56^Fe radiation fluence (data points at 1.1 and 3 particles/100μm^2^). **C**. Representative images showing 53BP1^+^ DNA double stranded break repair foci in the same mouse strains at 4 h and 24 h post-irradiation with 600 MeV/n ^56^Fe particles at the fluence of 3 particles/100μm^2^ and in sham-irradiated control.

We used two radiation response-relevant phenotypes for GWAS (**Fig. 1B**.) The first is Foci Per Gray (FPG), a quantification of the irradiation response using the average number of 53BP1^+^DNA repair foci per nucleus, per radiation dose (in Gy), measured at each time point for each radiation quality and mouse strain. FPG numbers followed previously reported kinetics of increasing at early time points (4 h and 8 h) post-irradiation that reflects ionizing radiation-induced DNA damage, followed by a gradual decrease at later time points (24 h and 48 h) due to ongoing DNA repair (Pariset *et al*. 2020b). FPG differences between strains for early time points also reflect the amount of double-strand break DSB clustering into DNA repair domains, a phenotype critical for high-LET radiation risk (Neumaier *et al*.; Vadhavkar *et al.*), since increased clustering is associated with lower repair and worse physiological outcomes. Representative images from *ex vivo* irradiated fibroblasts from two mouse strains with low and high FPG, CC019 and CBA, respectively, are shown in **Fig. 1C** at 4 h and 24 h after 600 MeV/n ^56^Fe irradiation and in sham conditions.

The second measured phenotype is Background (BGD), which is the average number of 53BP1^+^ foci at sham irradiation conditions, and reflects the amount of spontaneous DSBs. BGD was averaged for each strain across all irradiation experiments and all time points, after excluding statistical outliers (>2 standard deviations above or below the mean for the strain). Notably, although both male and female mice were used as a source of *ex vivo* cells, we did not observe sexual dimorphism in any FPG or BGD values.

To identify SNPs associated with FPG and BGD phenotypes, we performed a genome-wide association study (GWAS) based on sequencing the five inbred strains and known genotypes from the ten CC strains, accounting for kinship (described in **Materials and Methods**). **Fig. 2** shows representative Manhattan plots depicting the *p* values of the associations between SNPs and FPG 4 hours post-irradiation with X-rays, 350 MeV/n ^40^Ar and 600 MeV/n ^56^Fe, as well as background DNA repair foci. All SNP-phenotype associations are listed in **Suppl. Table 1**.

**Figure 2.**
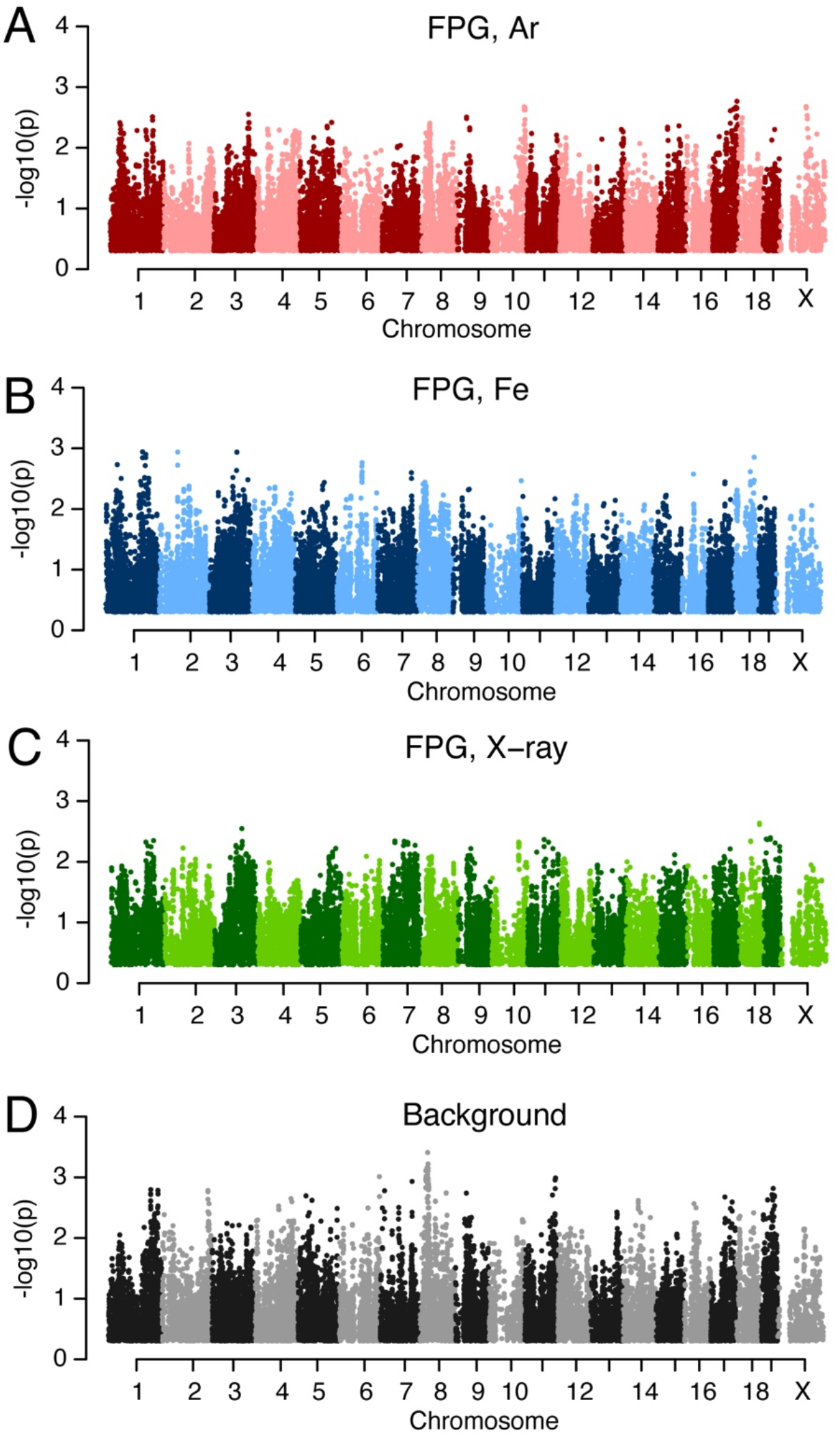
SNPs associated with DNA repair after irradiation and spontaneous DNA damage. Quantification was performed 4 h post-irradiation by X-rays and high-LET particles (^40^Ar, ^56^Fe). X-axis, chromosome. Y-axis, −log_10_(*p*), *p* values converted from *qtl2* LOD score.

### Genes and pathways associated with background DNA repair and responses to ionizing radiation

We characterized the quantitative trait loci (QTLs) linked with the SNPs identified by our GWAS analysis. SNPs were mapped to a gene if they fell within 25,000 base pairs of the gene. The resulting set of genes was analyzed using an additive approach of the Fisher method and addCLT to identify the genes with FDR-adjusted *p* values below 0.05, which were then mapped to pathways using the Reactome database. Significantly enriched pathways were identified using FGSEA and ORA test, and p values from each method were combined using addCLT to produce a single additive *p* value per pathway (see **Suppl. Figure 1** and **Materials and Methods** for a more detailed explanation). The key biologically significant genes and pathways significantly associated with FPG or BGD phenotypes are summarized in **Figure 3**, while the p-values for each gene and pathway are available in **Suppl. Tables 2-9**).

**Figure 3.**
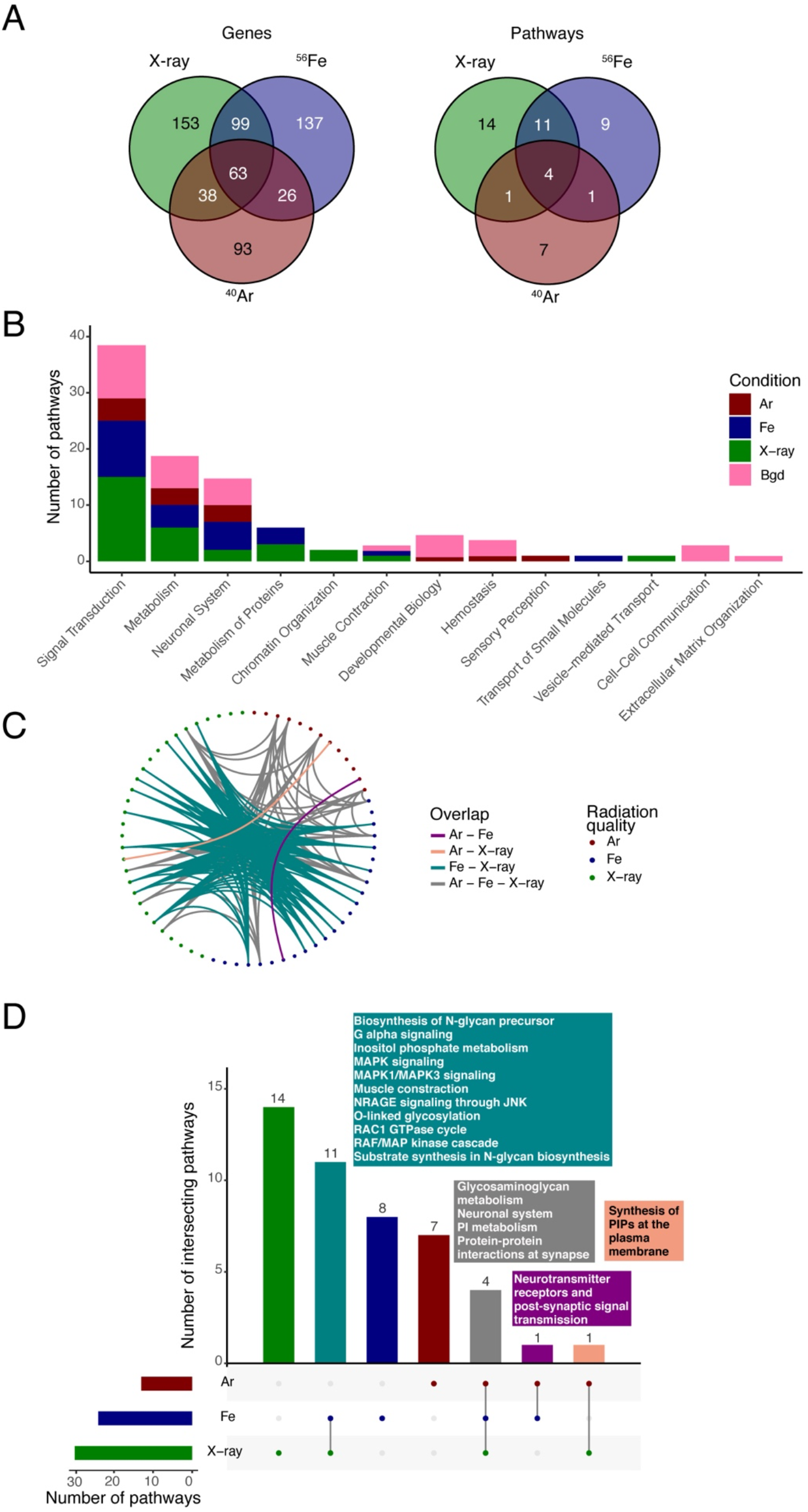
SNP, gene and pathway associations with DNA damage among radiation qualities. **A, B**.Diagrams of overlapping pathways among different radiation qualities. **C**.Bar graph of significantly enriched pathways in response to different radiation qualities and time points. Major pathway groups are listed on the X axis. **D**.UpSet diagram of overlapping pathways among radiation qualities across all time points. Pathways that overlap among multiple radiation qualities are listed in text insets shaded with matching color to the bars (teal: X-ray and 600 MeV/n ^56^Fe, orange: 350 MeV/n ^40^Ar and X-ray, purple: 350 MeV/n ^40^Ar and 600 MeV/n ^56^Fe, grey: all three radiation qualities). Radiation qualities: X-rays, 350 MeV/n ^40^Ar and 600 MeV/n ^56^Fe.

Some of the same genes and pathways were associated with both spontaneous and radiation-induced DNA repair (i.e. both BGD and FPG phenotypes). Out of 63 genes that were significantly associated with FPG induced by all three radiation qualities, 44 genes were also linked with BGD, while 19 genes were specific to FPG and 343 genes specific to BGD. This high overlap is consistent with our previous results that spontaneous DNA repair can be used as a biomarker of radiation sensitivity observed in radiotherapy patients (Pariset *et al*. 2020a), just like FPG is a biomarker for radiation sensitivity in mice *in vivo* (Pariset *et al*. 2020b). In addition, the high number of genes associated with BGD compared to FPG phenotype indicates that background DNA repair is determined by a wide variety of genetic factors, while DNA repair in response to an environmental stressor is regulated by comparatively few genes.

The shared genes that were associated with both FPG and BGD primarily fell into pathways associated with cell signal transduction and metabolism, as well as the nervous system (**Fig. 3B**). Meanwhile, the pathways linked with transport and chromatin organization were specific to radiation responses (FPG) and not spontaneous DNA damage (BGD), likely influenced by the fact that ionizing radiation leads to chromatin remodeling (Costes *et al*. 2010). On the other hand, cellular development and hemostasis as well as extracellular matrix organization (both of which are clusters of pathways involved in immune/hematopoietic responses) were linked with background (BGD) DNA repair.

### Comparing early and late DNA damage responses

In addition to quantifying the genes and pathways that were significantly associated with DNA repair across all time points, we split the time points into early (4 and 8 h after irradiation) and late (24 and 48 h after irradiation), representing initial DNA repair and persistent DNA damage and repair respectively.

The genes and pathways associated with early and late responses (**Fig. 4** and **Suppl. Tables 4-9**) partially overlapped: out of the 56 genes shared between all radiation qualities early and 94 late, 24 overlapped (**Fig. 4A**) with the main functions including DNA damage and tumorigenesis (*Cables1*, *Chst9*, *Dpysl3*, *Prkcb*, *Prkn*), the nervous system (*Dpysl3*, *Gls*, *Kcnn2*, *Mcph1*, *Ncald*, *Prkn*) and the immune/hematopoietic systems (*Flt3*, *Fut10*, *Il1rapl2*, *Rab7b*, *Stat4*, *Taf4b*, *Vps45*). Thus, these genes may be described as associated with a persistent phenotype of ionizing radiation.

**Figure 4.**
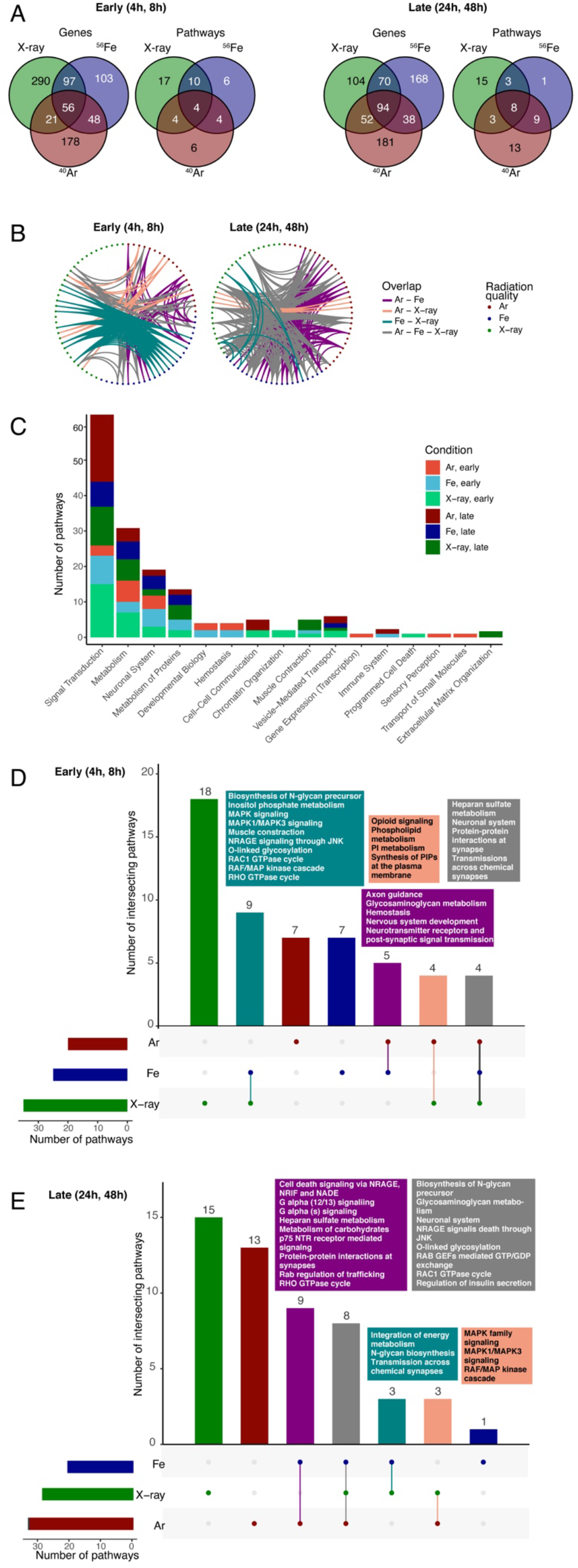
SNP, gene and pathway associations with DNA damage at different time points after irradiation. **A, B**. Diagrams of overlapping pathways among different radiation qualities. **C**. Bar graph of significantly enriched pathways in response to different radiation qualities and time points. Major pathway groups are listed on the X axis. **D, E**. UpSet diagram of overlapping pathways among radiation qualities early (**D**, 4 – 8 h) and late (**E**,24 – 48 h) after irradiation. Pathways that overlap among multiple radiation qualities are listed in text insets shaded with matching color to the bars (teal: X-ray and 600 MeV/n ^56^Fe, orange: 350 MeV/n ^40^Ar and X-ray, purple: 350 MeV/n ^40^Ar and 600 MeV/n ^56^Fe, grey: all three radiation qualities). Radiation qualities: X-rays, 350 MeV/n ^40^Ar and 600 MeV/n ^56^Fe.

Out of the 4 pathways shared between all radiation qualities early and 8 late, only 1 (“Neuronal system”) overlapped (**Fig. 4A, Suppl. Tables 8-9**). In addition, both time points also involved either glycosaminoglycan (late) or heparan sulfate (a specific type of glycosaminoglycan; early) metabolism. Glycosaminoglycans are expressed on most mammalian cells and are involved in cell signaling during carcinogenesis, including in the nervous system, as well as inflammation (Afratis *et al*. 2012; Morla 2019). Out of the non-overlapping pathways and genes, early responses primarily involved nervous system functions, while late response pathways were involved in cell signaling and cellular metabolism.

### Comparing responses to high and low LET radiation

Comparing the genes and pathways associated with FPG responses across all irradiation qualities, the strongest overlap was later after irradiation (**Fig. 4A**): 94 genes / 8 pathways compared to 56 genes / 4 pathways, indicating that radiation quality is more likely to affect initial, but not persistent DNA damage.

At early time points after irradiation the strongest overlap was between 600 MeV/n ^56^Fe and X-rays, possibly associated with the fact that these two qualities resulted in the highest doses: up to 0.82 Gy 600 MeV/n ^56^Fe, up to 4 Gy X-ray, compared to 0.5 Gy 350 MeV/n ^40^Ar. These pathways primarily involved cell signaling via MAP kinases, which have a wide range of pro-inflammatory and tumorigenic functions, as well as RHO GTPases (**Fig. 4 C, D**). RHO GTPases regulate cellular dynamics, including cell cycle and cellular migration (Hodge and Ridley 2016). They are involved in adverse cancer responses to therapeutic radiation, specifically, the formation of radiation-induced metastases (Zhai *et al*. 2006; Burrows *et al*. 2013), therefore are promising cellular targets for cancer treatment.

Other overlapping pathways between 600 MeV/n ^56^Fe and X-ray irradiation included cell death signaling via NRAGE, NRAGE plays a role in homologous recombination, which is the primary repair mechanism of double-stranded DNA breaks (Yang *et al*. 2016), and has a complex role in carcinogenesis: it suppresses metastasis, but also increases radiation resistance, which is harmful in the context of therapeutic radiation (though advantageous in the context of space radiation). In contrast, the pathways at early time points that were shared by high-LET particle, but *not* X-ray irradiation, were primarily focused on nervous system functions and glycosaminoglycan metabolism.

At later time points after irradiation we observed increased overlap in genes and pathways that were involved in high-LET particle radiation responses (600 MeV/n ^56^Fe and 350 MeV/n ^40^Ar), but were not shared with low-LET X-ray responses (**Fig. 4A, B**), likely due to the clustering and persistence of RIFs that are characteristic of high-LET particle irradiation (Penninckx *et al*.

2019). The high LET-specific pathways primarily regulate cell signaling via G alpha proteins and p75 receptors (**Fig. 4 C, E**), indicating them as suitable targets for high LET radioprotection and reduction of persistent DNA damage. Both G proteins and p75 signaling are associated with tumorigenesis (Johnston *et al*. 2007; Suzuki *et al*. 2009; Maziarz *et al*. 2020), and p75 is also involved in neurodegeneration (Knowles *et al*. 2009).

Finally, we uncovered multiple gene and pathway-level associations between radiation-induced DNA damage and the nervous system. Ionizing radiation is a major neurotoxic factor: high-LET particles have been shown to cause neuroinflammation, neurodegeneration and behavioral deficits in inbred C57BL/6J and transgenic Tg(Thy1-EGFP) MJrsJ mice (Parihar *et al*. 2016; Krukowski *et al*. 2021). However, our results suggest that different mouse strains might not be equally susceptible to ionizing radiation-induced neurotoxicity.

Most of the genes (e.g. *Dpysl, Prkn, Gls*) that were associated with *ex vivo* radiosensitivity have systemic functions in regulating oxidative stress, mitochondrial functions and carcinogenesis (Wang *et al*. 2010; Matsunuma *et al*. 2018; Kang *et al*. 2019), in addition to their involvement in neurodegeneration (Manivannan *et al*. 2013; Lynch *et al*. 2018; Yoshino *et al*. 2022). The variety of functions might explain why these genes were identified in fibroblast radiation responses *ex vivo* and suggests them as peripheral biomarkers of neurological risk for future experimental validation.

### Comparative radiosensitivity of 15 mouse strains

The majority of mouse studies on spaceflight and simulated space radiation have been conducted using only a few inbred strains, primarily C57BL/6 and BALB/c, or transgenic mouse lines with these strains as a background. However, our results indicate major differences in radiosensitivity between mouse strains and suggest deliberately selecting them based on experimental goals instead.

A representative comparison between mouse strains is presented in **Figure 5**. In this figure, all 15 strains are ranked based on the phenotype of interest, which is selected to be the average FPG at all time points and in response to all radiation qualities. In addition, the SNPs that mapped to selected carcinogenic and neurodegenerative genes (*Mcph*, *Ncald*, *Kcnn2*, *Gls*, *Dpysl3*, *Prkn*, *Cables1*, *Chst9*, *Prkcb*) that were significantly associated with FPG in the same conditions (i.e. shared by all time points and all radiation qualities) are listed for each mouse strain. The nucleotides are colored blue to match the one found in the most radioresistant strain (CC019) and yellow to match the most radiosensitive strain (CBA).

**Figure 5.**
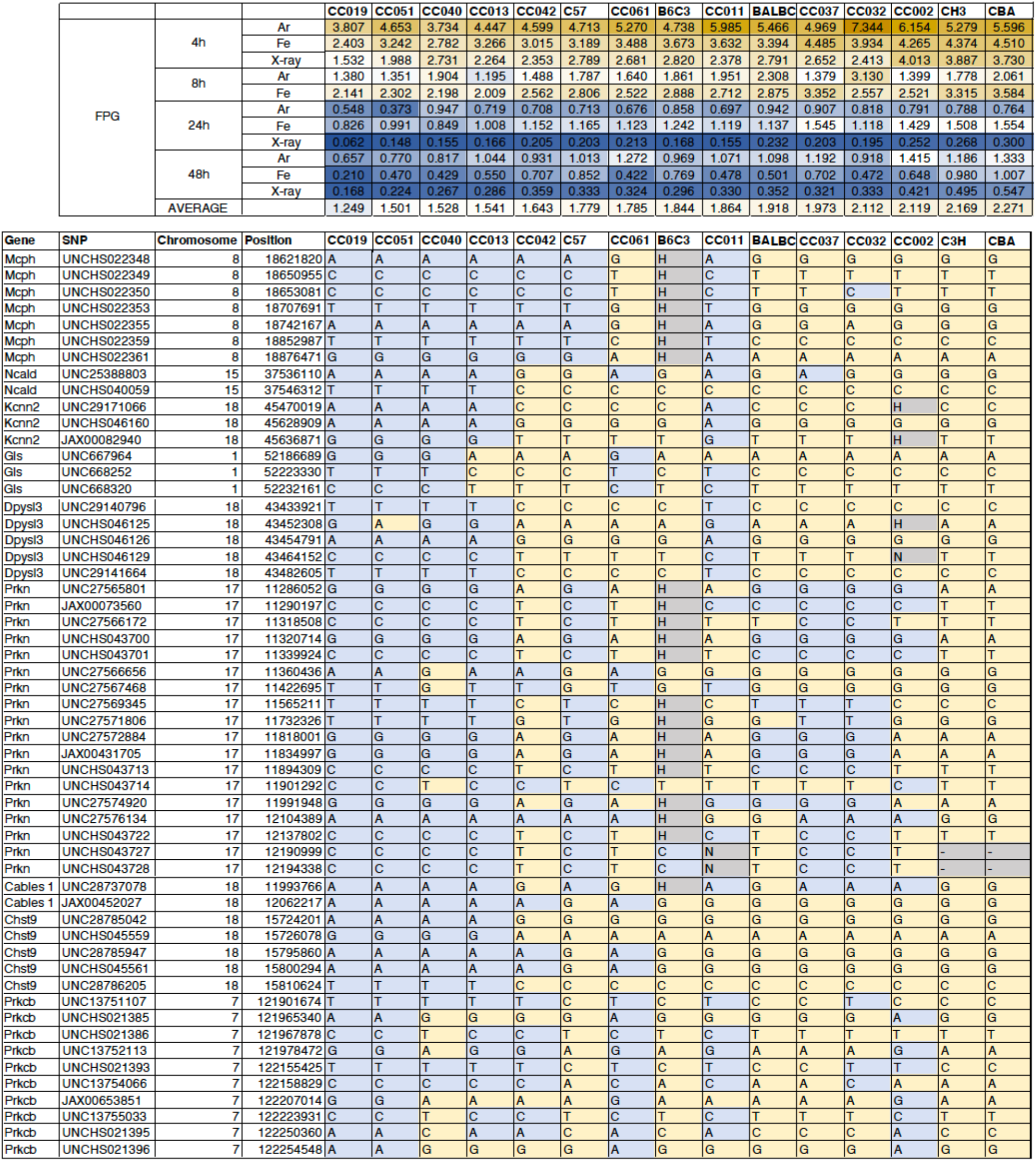
A representative example of comparative radiosensitivity of 15 mouse strains. Top table, FPG in response to different radiation qualities and time points. Strains are ranked by average FPG. Bottom table, SNPs mapped to selected genes with neurological and carcinogenic functions that were significantly associated with FPG across all radiation qualities and time points. Only SNPs with different nucleotides in the most radiosensitive and the most radioresistant strains listed. Blue, nucleotides matching the most radioresistant strain (CC019). Yellow, nucleotides matching the most radiosensitive strain (CBA).

This representative example shows the wide variability between mouse strains both in radiation-induced DNA damage responses and in the genotypes associated with it. Specifically, multiple CC strains (CC019, CC051 CC040, CC013) are comparatively radioresistant, while CC002, CH3 and CBA are particularly radiosensitive. Thus, using a combination of these strains in an experiment would be beneficial by covering a broader range of radiation responses, in this way reducing the probability of a response that does not reach a threshold or the opposite confounding factor of a response that is too strong and saturates the measurement. Furthermore, individual mouse strains might be selected based on interest in a specific gene or a group of genes with shared functions (e.g. neurodegeneration) or a particular radiation quality, dose or timepoint.

Some of our observed mouse strain differences in *ex vivo* radiosensitivity have been previously reported *in vivo* as well. For example, BALB/c has higher *ex vivo* radiosensitivity than C57BL/6 in our study, and BALB/c mice are also known to be more susceptible to radiation-induced cancers *in vivo* than C57BL/6 (Storer *et al*. 1988). However, when assessed within the entire range of the 15 mouse strains used in this investigation, C57BL/6 and BALB/c are much more phenotypically and genotypically similar to each other than some of the CC strains, suggesting CC as a better model to understand relative cancer susceptibility.

Although *in vivo* studies on CC mouse strains have been limited, a similarly high variability between strains has been observed in a transgenic melanoma model (Ferguson *et al*. 2015), where the CC19 mouse strain, which was particularly radioresistant in our study, consistently showed little melanoma progression. On the other hand, *spontaneous* tumor development in non-transgenic CC mice (Wang *et al*. 2019) was not associated with our results of either *ex vivo* radiosensitivity or spontaneous (background) *ex vivo* DNA damage, indicating its limitations as a biomarker. In addition, a study on the cardiotoxic side effects of doxorubicin, a DNA double strand-break inducing agent used in chemotherapy, showed that some of the CC strains that developed the most doxorubicin cardiotoxicity *in vivo* were among the most radioresistant *in vitro* in our study (Zeiss *et al*. 2019), suggesting separate genomic associations with DNA damage and with tissue degeneration.

<u>In conclusion</u>, we performed a genome-wide association study of *ex vivo* DNA damage responses to ionizing radiation in fibroblasts isolated from 15 collaborative cross and inbred mouse strains. We identified multiple SNPs, genes and pathways associated with spontaneous DNA damage and DNA damage responses to simulated deep space radiation (350 MeV/n ^40^Ar and 600 MeV/n ^56^Fe particles) and X-rays, which may underlie the differences in susceptibility to ionizing radiation among mouse strains. The genes and pathways were primarily linked to cellular signaling and metabolism as well as neurological impairments, and indicated different signaling cascades as suitable targets for limiting particle and X-ray radiation-induced DNA damage. Overall, this work shows how cell culture of different individuals can be used with the 53BP1^+^ radiation-induced foci assay to identify genes associated to radiation sensitivity. In light of the biomarker study we previously published on human blood cells exposed to radiation *ex vivo* (*Pariset et al. 2020a*), we suggest more GWAS studies should be conducted with similar assay to directly address this question in humans.

Our work serves as the first step in identifying the genes and pathways, which, following validation, will expand the application of genomic analysis to evaluate potential risk outcomes and therapeutic targets of ionizing radiation exposure during deep space travel. The availability of all 53BP1 expression-based DNA damage data for each *ex vivo* culture, radiation condition and time point combined with the genomics data in the NASA GeneLab -omics database provides a new tool for our community to further dive into radiation-relevant genotype-phenotype associations. More generally, our results and our publicly accessible data could be used to select mouse strains based on their radiation-relevant genomic characteristics for space radiation studies and countermeasure development.

## Data and code availability

All phenotypic and genomic data from this study are stored in the NASA GeneLab repository under GLDS-366, https://genelab-data.ndc.nasa.gov/genelab/accession/GLDS-366/ All code has been added to Supplementary Materials for reviewer access, and will be available on Github (https://github.com/duct317/Space-Radiation-Mouse-GWAS) upon publication. Raw image files are available upon request.

## Acknowledgments

We would like to thank Dr. A. Snijders and Dr. J.H. Mao at the Lawrence Berkeley National Laboratory for providing tissue from their Collaborative Cross cohort of mice.

## Figures and Figure Legends

**Supplementary Figure 1.**
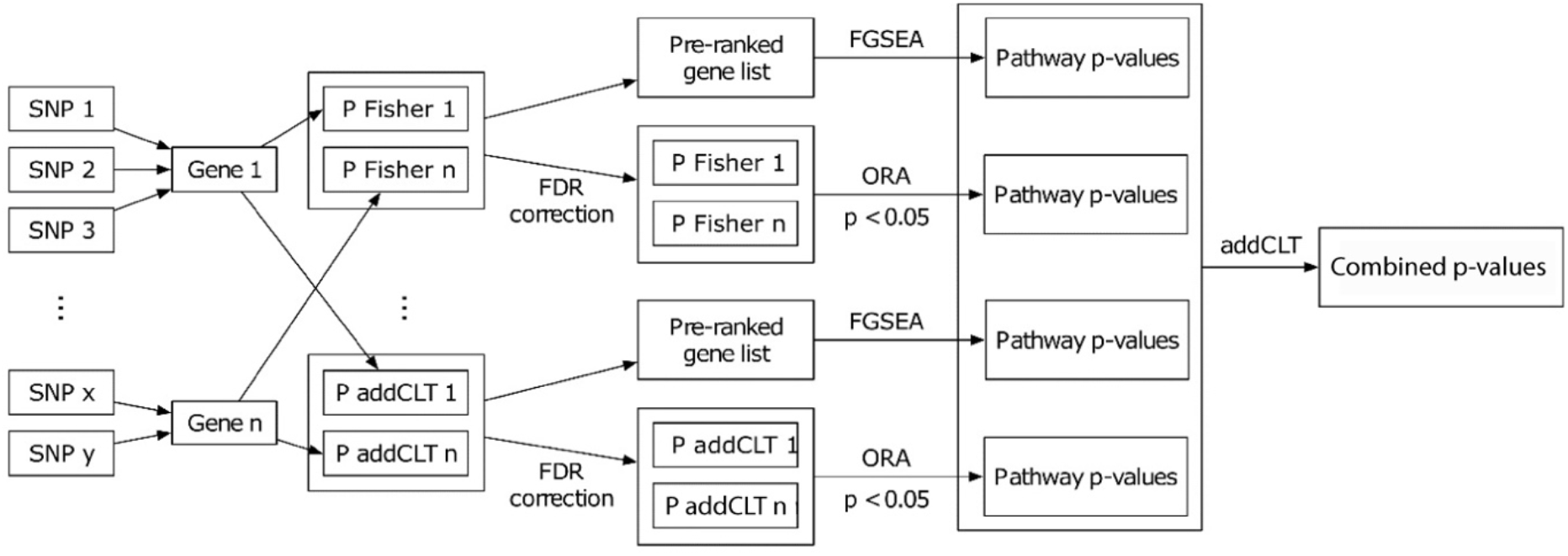
Gene and pathway analysis pipeline.

